# Male bumblebees adapt foraging to environmental conditions to sustain mate-seeking efforts

**DOI:** 10.1101/2025.09.08.674615

**Authors:** Natacha Rossi, Charlotte Doussot, Joseph L. Woodgate, Mathieu Lihoreau, Lars Chittka

## Abstract

Male bees navigate complex trade-offs between energy acquisition and reproductive signalling, yet their movement strategies remain understudied. Unlike workers that optimise foraging to support the colony, males forage independently to collect nectar and deposit sex pheromones on selected plants. Using high-resolution 3D tracking in an indoor flight cage, we investigated how the spatial arrangement of nectar and scent-marking sites, along with nectar availability, influence male movement patterns. We manipulated the distribution of feeders (artificial flowers) and scent-marking locations (branches), and varied nectar delivery rates, to assess effects on foraging, scent-marking, and patrolling. Males responded strongly to spatial structure: in clumped arrays with evenly spaced resources, movements between consecutive visits were shorter and more localised, while in dispersed arrays with irregular spacing, transitions were longer and more variable. The combination of dispersed spacing and low nectar availability imposed the highest foraging demands, resulting in fewer feeding events and reduced total feeding time. Despite these increased costs, males maintained consistent investment in reproductive behaviours, suggesting a prioritisation of mate-seeking over energy gain. Rather than reducing signalling, males adjusted their foraging strategy—favouring fewer but prolonged feeding bouts when nectar availability allowed. These findings reveal a unidirectional behavioural adjustment, in which foraging is modulated to sustain reproductive effort, and show how spatial resource structure and nectar availability together shape movement decisions in male pollinators.

**LAY SUMMARY:** Male bumblebees keep searching for mates even when food is scarce, adjusting how they forage to conserve energy. Using 3D tracking in a controlled environment, we found that males reduce foraging effort but maintain reproductive behaviours. This shows how animals balance survival and reproduction in changing environments, offering new insights into pollinator behaviour and resilience.

## INTRODUCTION

Across taxa, animals adjust how they allocate time and energy – for example, balancing foraging with reproductive behaviours – based on both internal conditions (such as energy reserves) and external cues (such as resource availability, competition and predation risk) (Dougherty, 2021; Gavassa et al., 2013; Westwood et al., 2025). For males in particular, behavioural decisions, such as when and where to forage or engage in reproductive signalling, often reflect trade-offs between energy acquisition and mating effort. When resources are scarce, males may prioritise energy acquisition. However, when resources are abundant or competition for mates is high, they may shift their investment towards mate-seeking (Billings et al., 2019; Fernlund Isaksson et al., 2022; Höbel, 2015; Meuche and Grafe, 2009; Wilde et al., 2023).

Movement strategies thus often reflect the functional demands of a given behaviour, whether it be foraging, defending resources, or seeking mates. For instance, hummingbirds defend fixed feeding territories to ensure exclusive access to high-reward resources that can be monopolised (Sargent et al., 2021). In contrast, butterflies move more randomly when searching for mates, a strategy suited to locating widely dispersed and unpredictable mates in open environments (Reynolds, 2006). Many nectar-feeding animals, such as bees and certain bats, establish stable, repeatable routes between feeding locations (traplines), which maximise energy efficiency and minimise travel costs in environments where resources are spatially stable but not defensible (Berger-Tal and Bar-David, 2015; Mailly et al., 2025).

Bumblebee workers (*Bombus* spp.) exemplify route optimisation: they learn sequences of flower visits that they adjust in response to changes in spatial layout and nectar availability (Cartar and Real, 1997; Chittka et al., 1997; Lihoreau et al., 2012; Ohashi and Thomson, 2009; Ohashi et al., 2007; Raine and Chittka, 2007; Thomson et al., 1997; Woodgate et al., 2017). However, male bumblebees follow a fundamentally different life history trajectory. Soon after eclosion from their pupae, males typically leave the nest permanently, no longer contributing to colony tasks such as foraging or brood care (Free, 1982; Kearns and Thomson, 2001). Instead, they roam independently, balancing the need to forage for themselves with the imperative to locate and attract mates (Frank, 1941; Freeman, 1968; Haas, 1946; Haas, 1949).

A key component of male reproductive behaviour in many *Bombus* species involves scent-marking patrol circuits during which they deposit pheromones on substrates such as branches, leaves, or stems as they establish regular flight paths to attract receptive queens (Frank, 1941; Haas, 1946; Haas, 1949; Haas, 1952; Harano et al., 2018; Kullenberg, 1973), a behaviour first described by Darwin (Freeman, 1968). These patrol routes are typically stable, with males revisiting specific marking sites repeatedly (Frank, 1941; Harano et al., 2018). These circuits share some structural resemblance to the traplines used by workers foraging for nectar (Thomson et al., 1997), but may reflect different behavioural priorities and decision-making rules.

Although bee males contribute to pollination and influence mating dynamics (Kraus et al., 2009; Ogilvie and Thomson, 2015), their movement strategies remain poorly understood. In contrast to workers, whose foraging patterns and spatial behaviours have been extensively studied, male movement ecology—particularly how it is shaped by environmental factors such as resource distribution and nectar availability—has received limited attention. Quantifying male foraging and reproductive behaviours in controlled spatial environments is therefore essential to understand how they navigate ecological trade-offs and to clarify their role in pollination systems.

To address this gap, we used high-resolution 3D video tracking of the spatial movement patterns of male *Bombus terrestris* in a controlled flight cage environment. This allowed us to track individual males with unprecedented accuracy, enabling detailed analysis of their movement decisions over extended periods. The flight cage contained artificial flowers (providing nectar) and scent-marking locations (non-rewarding branches), mimicking the dual-resource environment males encounter in nature. We systematically manipulated the spatial arrangement of these elements and the nectar delivery rate to assess how males navigate trade-offs between feeding and reproductive efforts. Specifically, we sought to answer four key questions: (1) Do males trade off foraging effort for reproductive signalling when nectar availability or spatial resource configuration changes? (2) What movement strategies do males follow when visiting successive items? (3) Do they adjust the frequency of revisits over time? (4) Do they establish repeatable routes, and if so, are these routes affected by spatial configuration and nectar reward rates?

## METHODS

### Experimental setup

Experiments were conducted in a flight cage (300 cm L × 300 cm W × 150 cm H) within a controlled laboratory environment between June 2021 and July 2022. The cage was enclosed with white tulle fabric (Handi Stitch, Amazon UK), a fine mesh material with an estimated pore size of c.a. 0.5mm, suitable for containing bumblebees while allowing airflow and visibility. The cage was surrounded by white curtains to standardise visual cues for the bees and to optimise 3D tracking by providing strong contrast between the bees and the background. The floor was covered with laminated printouts of red and white Julesz patterns, which are random-dot textures used to provide high spatial frequency contrast without distinct landmarks (Julesz, 1962). These patterns enhanced optic flow cues for flight control and improved the accuracy of 3D tracking by providing a high-contrast background against which bees could be reliably detected. A 40 × 41 cm piece of purple-painted cardboard was placed between the camera bases to serve as a spatial reference point for the bees. Environmental conditions were maintained at 21.8 ± 0.02 °C and 46.2 ± 0.3% relative humidity, recorded continuously using a digital sensor.

### Subjects

We used male *B. terrestris* from ten commercial colonies (each containing a queen and 40–50 workers) and from additional male-only boxes (∼ 50 males per box), all purchased from Agralan Ltd. (Wiltshire, UK). Males in the boxes were almost certainly sourced from multiple colonies as a single queen typically produces only a small number of males. Colonies were housed in wooden nest boxes (28 × 16 × 11 cm) lined with cat litter to regulate moisture levels. A 25% (w/w) sucrose solution was provided via feeders, and nests were provided with Natupol pollen (Koppert, Suffolk, UK) twice weekly.

A total of 62 males were initially included in the experiment. However, 11 were excluded because of escape, persistent flight instability (e.g. repeated falling), feeder dislodging, incomplete video recordings, or tracking errors, resulting in a final sample size of 51 males.

### Flight cage items

To simulate a realistic environment for foraging, scent-marking, and patrolling, we arranged six *Cotoneaster horizontalis* branches and six artificial flowers in the flight cage. *C. horizontalis* is a shrub whose flowers are naturally visited by bumblebees (Corbet and Westgarth-Smith, 1992) and was used as a substrate for male scent-marking behaviour. Branches were selected to have minimal lateral branching and measured approximately 15 cm, providing a relatively uniform and accessible structure for landing and marking. Each branch was mounted on a 3D-printed white tripod (10 cm high) to ensure stability and consistent presentation across trials.

The artificial flowers (Figure 1a) were designed to deliver controlled amounts of nectar substitute (BIOGLUC®, Biobest, Belgium), which was used instead of sugar water as it does not crystallise as quickly, an important consideration given the extended duration of our experiments. Each flower consisted of a white 3D-printed tripod (5 cm high), supporting a 6 cm wooden cylinder (1 cm radius) with tubing (3 mm outer diameter, 1 mm inner diameter) to facilitate nectar flow. A yellow square plastic chip (2.5 × 2.5 × 0.5 cm) served as a visual cue, and an Eppendorf tube tip (0.7 cm deep) held the sucrose solution. The Eppendorf tube depth was selected to match the tongue length of *B. terrestris* males (c.a. 0.7cm, Gérard et al., 2023), ensuring efficient feeding.

**Figure 1.**
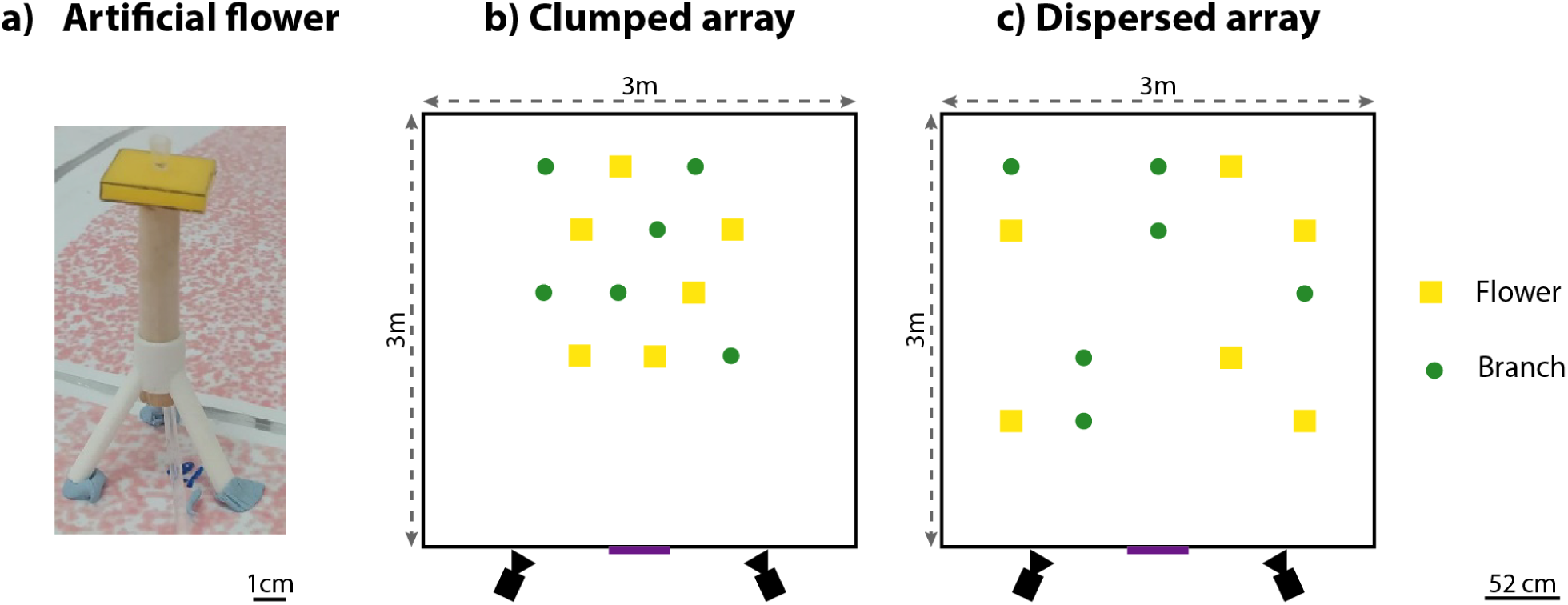
Experimental spatial arrangements and artificial flower. (a) Artificial flower: Photograph of an artificial flower, showing the white 3D-printed tripod, vertical wooden support, yellow plastic visual cue, and sucrose feeder tip. (b) Clumped array: Branches (green circles) and artificial flowers (yellow squares) were arranged in a uniform grid at 52 cm intervals within a 135 x 135 cm area (c.a. 1.82 m^2^), simulating a densely packed patch. (c) Dispersed array: Items were placed at irregular distances across a 208 x 208 cm arena (c.a. 4.33 m^2^), using a 5 x 5 grid as a placement guide. Nearest-neighbour distances ranged from 52 cm to c.a. 214 cm, resulting in a more spatially heterogeneous layout. 3D tracking cameras are represented in black and the purple bar represents a central landmark. The flight cage measured 3 x 3m.

Artificial flowers were automatically refilled every hour using Tempatron AM24 drive motors and a gear kit-controlled cam sequencer system (RS Components, Corby, UK). The refill system was active for 15 minutes at the start of each hour, followed by a 45-minute pause, repeating this cycle continuously throughout the day. Two different sucrose delivery rates were implemented to simulate conditions of high and low nectar availability. Under the high-rate condition, nectar was delivered at 4.64 ± 0.16 μl/min, resulting in 69.6 ± 0.16 μl per flower over each 15-minute refill period. In contrast, under the low-rate condition, nectar was dispensed at 1.63 ± 0.23 μl/min, yielding 24.4 ± 0.23 μl per flower per refill cycle. While these values exceed typical nectar secretion rates of individual flowers in bumblebee-pollinated species such as *C. horizontalis* (c.a. 0.01-0.05 μl/min; Dmitruk et al., 2022) and *Digitalis purpurea* (c.a. 0.04 μl/min; Gaffal et al., 1998), they are comparable to the upper range observed in highly productive species like *Lamium maculatum* under continuous foraging conditions (c.a. 3.7 μl/min; Marina et al., 2004).

Given that the mean maximum crop fill in *B. terrestris* males is 118.5 ± 45.8 μl (N=20; Wolf and Chittka, 2016), bees would need to collect all nectar available in two fully replenished high-rate flowers or five fully replenished low-rate flowers to fill their crop. This setup thus allowed us to investigate how changes in nectar availability influenced male bee foraging behaviour and movement patterns.

### Spatial arrangements of items

Two spatial configurations were tested, representing contrasting resource distributions: a clumped array and a dispersed array (Figure 1b-c).

In the clumped array, items (branches and artificial flowers) were arranged at regular intervals (52 cm) within a 135 cm x 135 cm area (c.a. 1.82 m^2^), creating a dense, uniformly spaced layout that mimicked a tightly packed patch of vegetation.

In the dispersed array, items were placed at irregular distances across a 208 cm x 208 cm area (c.a. 4.33 m^2^), guided by a 5 × 5 placement grid to achieve a heterogenous layout with variable spacing between items. This resulted in a more spatially complex environment, with nearest-neighbour distances ranging from 52 cm to c.a. 214 cm and fewer local neighbours per item.

Note that the total surface area differed between configurations, and spatial regularity was not controlled independently of area size. Consequently, any behavioural differences observed between the two spatial arrangements reflect the combined effects of spacing structure and density.

A total of 51 males were tested across the spatial and nectar availability conditions: 12 in the dispersed array with low nectar availability, 12 in the clumped array with low nectar, 12 in the clumped array with high nectar, and 15 in the dispersed array with high nectar.

### Training procedure

To facilitate learning, six males were removed from their colony or male box one day before testing and marked with numbered, coloured disks (Bienen-Voigt & Warnholz GmbH & Co. KG, DE) glued to their thoraces for individual identification. The bees were then placed in a wooden training box (28 × 16 × 11 cm) containing six yellow plastic chips (matching those in the artificial flowers). A drop of BIOGLUC® was manually pipetted onto each cuboid to establish a colour-reward association. Refills were provided immediately after depletion, and the bees were observed until they had contacted the sucrose with their proboscis at least once.

### Testing procedure

Male bees were released into the flight cage at c.a. 15:30 on the day prior to testing. The cage operated under a light-dark regime of 8 hours of light and 16 hours of darkness, with lights on from c.a. 09:00 to 17:00. Following release, bees experienced c.a. 1.5 hours of light (from 15:30 to 17:00), during which they began initial exploration and familiarisation with the new environment. On the day of testing, males were observed until at least one individual had scent-marked a branch, fed from an artificial flower, and demonstrated sustained flight. Once a suitable bee was identified, individual testing began. Only one bee was tested at a time to eliminate social interference. All other bees were removed and returned to the training box. The focal bee was then recorded for six hours, from 09:30 to 15:30, using two Basler GenICam cameras for 3D tracking. This consistent 6-hour observational window allowed comparison across individuals and conditions while controlling for potential circadian variation.

### Data analysis

We used the software package Track3D (Noldus Information Technology, Wageningen, The Netherlands) as an add-on to EthoVision XT 16.0.1538 to generate three-dimensional tracking data. EthoVision XT extracted 2D coordinates from two synchronised Basler video cameras, and Track3D combined these into a set of 3D coordinates (X, Y, Z) using stereo triangulation. This reconstruction was based on prior calibration with CentroidFinder, which aligned the image coordinates to real-world spatial dimensions. The cameras recorded bee positions at 30 frames per second.

#### Behavioural classification

Behavioural modes were classified using a simple hand-crafted algorithm using a small number of thresholds for movement statistics. These thresholds were initially informed by exploratory analyses and comparison to expert-labelled segments, but the final classification rules were selected to approximate expert observations as objectively and consistently as possible. Three distinct behavioural modes were analysed: Feeding, Scent-Marking, and Patrolling.

First, the 3D positional data for each track were broken into 0.5s segments and movement statistics calculated for each segment: speed, turning angle, distance travelled, and proximity to the nearest flower or branch. Segments in which the mean movement speed exceeded 70mm/s were classified as flight. It was not possible to use movement speed to differentiate between walking and sitting still because bees often move very slowly. Thresholds for angular momentum and distance moved can be effective at differentiating different walking and sitting behaviours but since the behaviours we were interested in (feeding and scent-marking) can involve a range of movement speeds we treated all walking and sitting movement as the same for the purposes of further behavioural classification.

Feeding was defined as any instance in which a bee was recorded walking or sitting within 20 cm of a flower and no more than 5.5 cm above or below the flower platform. This spatial threshold was used because the resolution and distance of the cameras did not allow for direct visual confirmation of feeding, and slight positional variation of flower placement occurred between trials despite marked floor guides. The 20 cm radius accounts for this variability while ensuring a consistent, conservative estimate of flower visits likely associated with feeding. To ensure interactions were specific to the flower platform, body movement had to be predominantly horizontal, with a horizontal-to-vertical movement ratio exceeding 1.5.

Scent-Marking was characterised by a bee walking or sitting within a 20 cm radius of an item at a height of no more than 1 cm above it, where vertical body movement was predominant (horizontal-to-vertical movement ratio below 1. In the context of this study, bees exhibited such behaviour on both artificial flowers and branches.

We used the term Patrolling to describe flight behaviour near flowers or branches involving slow, highly manoeuvrable movement, consistent with environmental inspection in search of mates. A patrolling bout began when a bee entered a 20 cm radius around a flower or branch and flew no more than 10 cm above it, while maintaining a flight speed below 50 cm/s and an angular velocity (i.e., rate of change in heading direction) above 125°/s. These thresholds were chosen to distinguish patrolling from faster, straighter transit flights, capturing the characteristic tight turning and hovering-like motion typical of inspection behaviour. A patrolling event was considered to have ended when the bee: (1) landed on a branch, the cage floor, or netting; (2) ascended more than 45 cm above the level of the items (suggesting transition to exploratory flight); or (3) went more than 5 seconds without meeting the criteria for further inspections.

#### Hypothesis testing

All statistical analyses were conducted in R (v.4.4.2; R Core Team, 2024), using the lme4 (Bates et al., 2015), emmeans (Lenth, 2024), and performance (Lüdecke et al., 2021) packages. We used linear mixed-effects models (LMMs) to test how male behaviour varied in response to spatial resource distribution (clumped vs. dispersed), nectar availability (low vs. high), and behaviour type (feeding, scent-marking, patrolling). All models included Male ID as a random intercept. Significance of fixed effects was assessed using Type III Wald χ² tests (car package; Fox and Weisberg, 2019), and estimated marginal means were extracted using emmeans with Tukey correction for multiple comparisons. Where applicable, predicted values were back-transformed from log-transformed response variables. Model fit was assessed using marginal and conditional R² (performance package) and model selection was based on Akaike’s Information Criterion (AIC), with preference given to the simplest model with fewer degrees of freedom.

Behavioural frequency (log-transformed) was modelled as a function of behaviour type, spatial resource distribution, nectar availability, and their interaction (behaviour × spatial distribution). Marginal R² = 0.203; Conditional R² = 0.648.

Total behavioural duration (log-transformed) was modelled with fixed effects of behaviour type, nectar availability, spatial resource distribution, and the interactions behaviour × spatial distribution and nectar availability × spatial distribution. Marginal R² = 0.187; Conditional R² = 0.522.

Per-event duration (log-transformed) was modelled as a function of behaviour type, spatial resource distribution, nectar availability, and their interactions (behaviour × spatial distribution, nectar availability × spatial distribution). Due to the large sample size, post hoc comparisons used asymptotic approximations. Marginal R² = 0.489; Conditional R² = 0.523. A minimum duration threshold of 0.5 s was applied to exclude transient events unlikely to reflect sustained behavioural states.

To analyse transition patterns in movement, the distance and angle between two consecutive visits were examined across different nectar availability conditions and spatial arrays using Python (version 3.11.7). The Mann-Whitney U test was used to compare distributions, and Cohen’s d was computed as an effect size measure to quantify differences. Return cycles (i.e. number of items visited before the male returned to the same item) were modelled using a Negative Binomial distribution, incorporating Male ID as a random effect, with nectar availability and array configuration included as interaction terms. Immediate revisits were also modelled using a Negative Binomial distribution, including Male ID as a random effect, along with array configuration and behaviour type as interaction terms. Immediate revisitation events were counted when a male visited the same location consecutively (e.g., sequence A → A) with no intervening visits to other items.

To assess whether male bees used repeated routes between items, we calculated the Determinism Index (DET) following Ayers et al. (2015). This index quantifies sequence repetition, where values closer to 1 indicate highly deterministic movement patterns, while values near 0 indicate random movement. DET was calculated using sequences ranging from three to five items in length. To evaluate whether movement patterns resulted from structured decision-making rather than randomness, DET values (excluding redirect revisits) were compared to those generated by movement models simulating specific navigation rules within the spatial array. We implemented four movement models to compare against observed bee behaviour: Nearest Neighbour (NN): Bees moved to the closest item; Direct Neighbour (DN): Bees moved to nearby, unoccluded items within 2× the nearest-neighbour distance and outside a 25° masking angle, representing a broader but vision-constrained local choice; Low Turn (LT): Bees moved in directions that minimised turning angle, mimicking persistence in directionality; Random (R): Bees selected the next item at random, simulating non-directed exploration. These models were implemented from scratch in Python using geometric and probabilistic rules. Normality of DET values was assessed using Shapiro-Wilk tests. Differences among models (Nearest Neighbour, Direct Neighbour, Low Turn, Random, and Real Bees) were evaluated using Kruskal-Wallis tests, followed by Dunn’s post hoc tests with Bonferroni correction for multiple comparisons.

All data and analysis code will be archived in a public repository and made available upon publication, in accordance with *Behavioural Ecology*’s data policy.

## RESULTS

### Behaviours

Male bumblebee behaviour varied in response to the spatial distribution of resources, with additional effects of nectar availability observed in specific contexts. These effects were primarily evident in feeding behaviour, whereas reproductive signalling behaviours (patrolling and scent-marking) remained comparatively stable.

#### Behavioural frequencies (*Figure 2a*)

We examined whether male bumblebee behaviour differed in frequency across spatial resource distribution (clumped vs. dispersed), nectar availability (low vs. high), and behaviour type (feeding, scent-marking, patrolling). This analysis aimed to determine whether males altered the relative allocation to foraging versus reproductive behaviours depending on environmental context.

**Figure 2.**
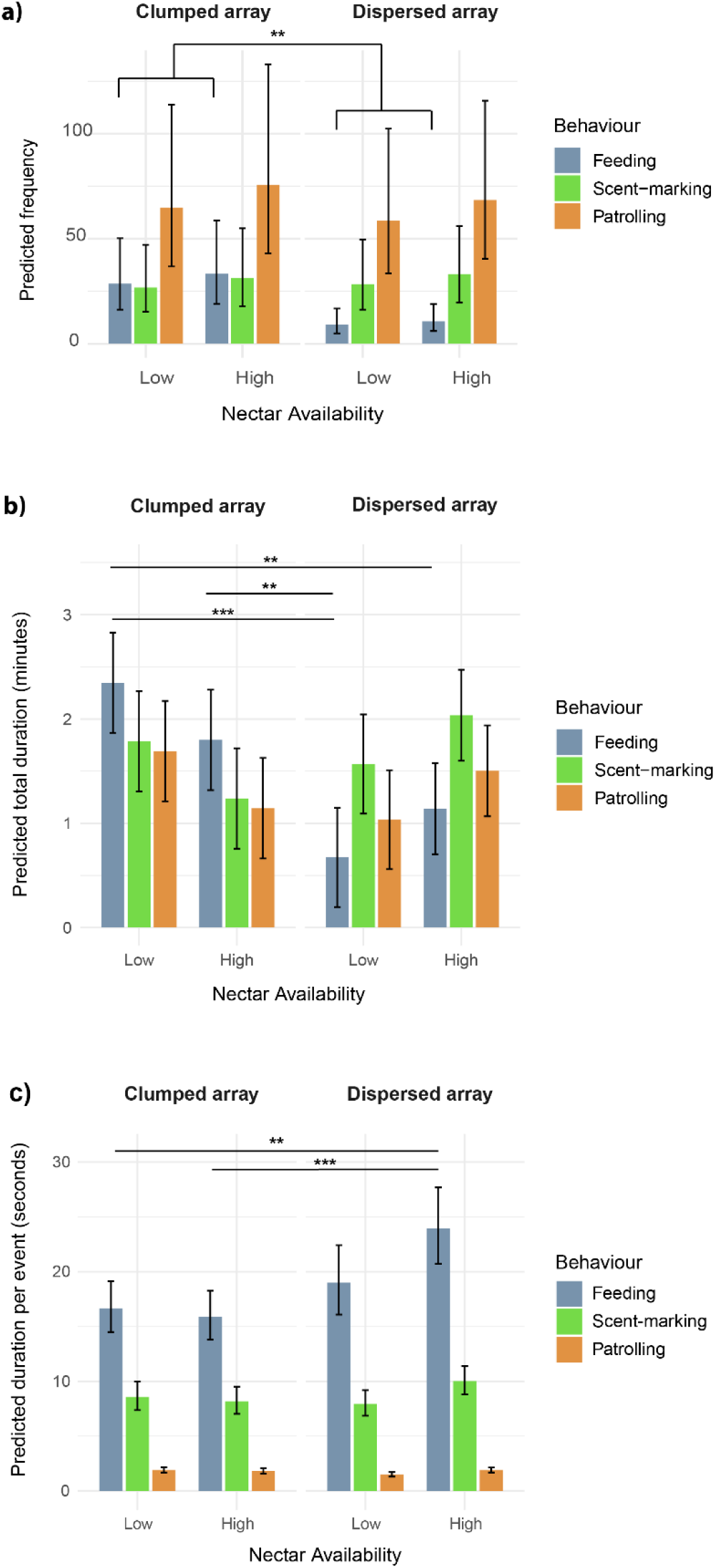
Predicted behavioural metrics for male bumblebees in relation to spatial resource distribution and nectar availability. (a) Predicted frequency of behaviours from a linear mixed model including behaviour, spatial distribution (clumped vs. dispersed), nectar availability (low vs. high), and their interaction. (b) Predicted total duration of each behaviour (in minutes). (c) Predicted per-event duration of each behaviour (in seconds). All values are back-transformed from log-transformed data. Error bars represent 95% confidence intervals based on estimated marginal means. All models included Male ID as a random effect. Treatments are shown across nectar availability (low vs. high) and spatial distribution (clumped vs. dispersed). Behaviour types are colour-coded and separated by facets in panels (a)–(c). A total of 52 males were tested: 13 in the dispersed–low nectar condition, 12 in clumped–low, 12 in clumped–high, and 15 in dispersed–high. Tukey-adjusted pairwise comparisons of estimated marginal means: ** *P* < 0.01, *** *P* < 0.001.

A linear mixed model revealed a significant interaction between behaviour type and spatial resource distribution (χ²(2) = 14.13, *P* < 0.001), indicating that behavioural frequencies varied depending on spatial arrangement. Main effects of behaviour type (χ²(2) = 17.99, *P* < 0.001) and spatial resource distribution (χ²(1) = 10.04, *P* = 0.0015) were also significant, reflecting overall differences in behaviour frequency across conditions. Nectar availability had no significant effect (χ²(1) = 0.29, *P* = 0.589).

Tukey-adjusted post hoc comparisons of estimated marginal means revealed that in the clumped array, patrolling occurred more frequently than feeding (estimate = –0.818, SE = 0.232, *P* = 0.0019) and scent-marking (estimate = 0.883, SE = 0.232, *P* = 0.0007), while feeding and scent-marking did not differ (*P* = 0.9581). In the dispersed array, feeding was significantly less frequent than both patrolling (estimate = –1.857, SE = 0.248, *P* < 0.0001) and scent-marking (estimate = –1.131, SE = 0.248, *P* < 0.0001), and patrolling also exceeded scent-marking (estimate = 0.726, SE = 0.219, *P* = 0.0037).

Within-behaviour comparisons showed that feeding frequency was significantly reduced in the dispersed array compared to the clumped array (estimate = 1.139, SE = 0.360, *P* = 0.0021), while no spatial differences were observed for patrolling or scent-marking (*P* > 0.76).

#### Total behavioural duration (*Figure 2b*)

We tested whether the total time spent performing each behaviour varied with spatial resource distribution, nectar availability, and behaviour type. This analysis aimed to assess how environmental conditions shaped males’ overall investment in foraging and reproductive behaviours.

A linear mixed model revealed a significant interaction between behaviour type and spatial resource distribution (χ²(2) = 27.65, *P* < 0.001), as well as an interaction between nectar availability and spatial distribution (χ²(1) = 6.14, *P* = 0.013). Main effects of behaviour (χ²(2) = 11.74, *P* = 0.0028) and spatial distribution (χ²(1) = 4.08, *P* = 0.044) were also significant. Nectar availability had no significant main effect (χ²(1) = 3.40, *P* = 0.065).

Tukey-adjusted post hoc comparisons showed that for feeding, total duration was significantly lower in the dispersed–low condition compared to both clumped–low (*P* < 0.0001) and clumped–high (*P* = 0.0075), but did not differ significantly from the dispersed–high condition (*P* > 0.35).

For patrolling and scent-marking, no significant differences in total duration were detected between conditions (*P* > 0.22), indicating consistent investment in these behaviours regardless of environmental variation.

#### Per-event behavioural duration (*Figure 2c*)

We tested whether males adjusted the duration of individual behavioural events in response to changes in spatial resource distribution, nectar availability, and behaviour type. This analysis aimed to assess whether males modulated how they fed or signalled depending on environmental conditions.

A linear mixed model revealed significant effects of behaviour type (χ²(2) = 5501.16, *P* < 0.001), as well as interactions between behaviour and spatial resource distribution (χ²(2) = 43.52, *P* < 0.001) and between spatial distribution and nectar availability (χ²(1) = 4.38, *P* = 0.036). The main effects of nectar availability and spatial distribution were not significant (*P* > 0.23).

Tukey-adjusted post hoc comparisons showed that feeding events lasted significantly longer than both scent-marking and patrolling events in all treatment combinations (*P* < 0.0001). For feeding, per-event duration was longest in the dispersed–high condition, which was significantly longer than in both clumped–high (*P* = 0.0004) and clumped–low (*P* = 0.0021). However, feeding durations in the dispersed– high and dispersed–low conditions did not differ significantly (*P* = 0.36).

Per-event durations for scent-marking and patrolling did not differ significantly across conditions (*P* > 0.06), indicating that the temporal structure of reproductive signalling was stable across environmental treatments.

Collectively, these results show that the dispersed–low nectar condition imposed the highest foraging cost, as reflected by the lowest feeding frequency and total feeding duration. Males did not reduce investment in patrolling or scent-marking under any condition. Instead, they adjusted their foraging effort by modifying how they fed: when foraging opportunities were limited, males fed less frequently and for shorter total durations and extended per-event durations only when nectar availability allowed.

This pattern indicates that male bumblebees respond to challenging foraging conditions not by reallocating time from reproductive behaviours, but by adapting the structure of their foraging efforts, optimising intake efficiency when resources are spatially dispersed and costly to access.

### Movement strategies

To examine how nectar availability and spatial configuration influenced movement behaviour, we analysed pairwise transitions between consecutive item visits, focusing on the distance and angle of each movement. These transition metrics allowed us to quantify how bees adjusted their foraging strategy across the four experimental conditions (Figure 3).

**Figure 3.**
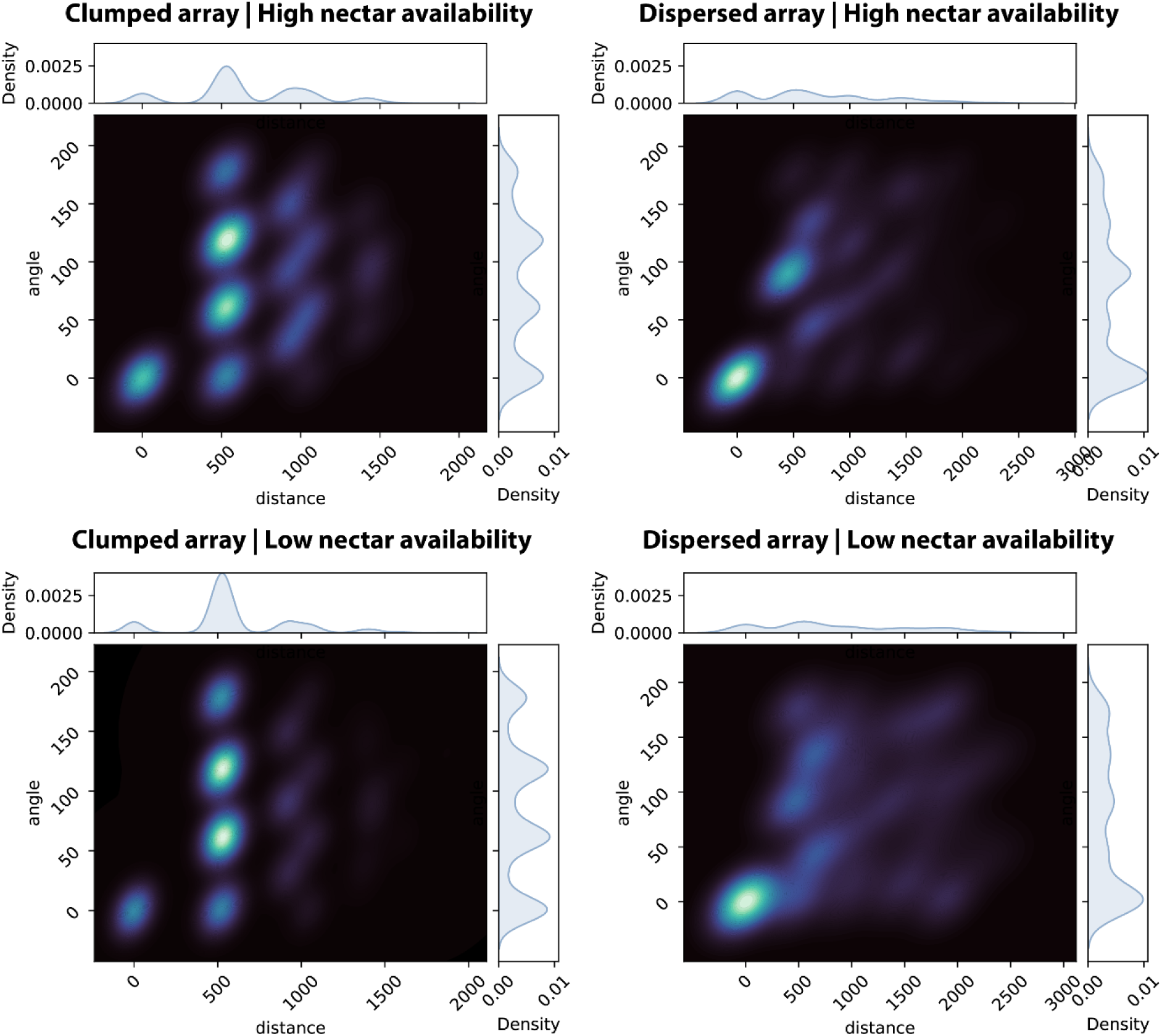
Density plots of movement transitions between consecutive visits to items (artificial flowers or branches), shown as a function of distance and direction (angle), across spatial arrays and nectar availability conditions. Each panel represents one of four experimental treatments: clumped (left) or dispersed (right) spatial arrays under high (top) or low (bottom) nectar availability. The x-axis indicates the distance (mm) between successive visits, and the y-axis shows the movement angle (°), relative to the arena’s x-axis (0° = forward, 90° = lateral, 180° = backward). Brighter areas represent higher densities of transitions. Marginal density plots show the distribution of distances and angles separately.

Across both spatial configurations, males tended to visit closer locations when nectar availability was low, consistent with a more localised foraging strategy under limited resources. In the clumped array, individuals moved significantly shorter distances at low nectar availability compared to high (U = 8,163,133, *p* < 0.001, *f²* = 0.18). A similar pattern was observed in the dispersed array, where males also reduced movement distances under low nectar availability (U = 2,259,686, *p* < 0.001, *f²* = 0.25). By contrast, when nectar was abundant, individuals travelled farther, likely reflecting a more exploratory strategy aimed at maximising intake from replenished sources.

Directionality of movement also varied with nectar availability, particularly in the dispersed array. Males exhibited more consistent directional movements when nectar availability was low (U = 2,490,456, *p* = 0.0186, *f²* = 0.05), although the effect size was small. This suggests that when resources are limited, males may return to previously visited locations rather than investing in broader exploratory searches.

Together, these findings indicate that males dynamically adjusted both foraging distance and directional consistency in response to resource abundance. When nectar was scarce, males adopted a more restricted movement style and prioritised known locations; when it was abundant, they broadened their range and explored more widely.

### Revisitation patterns and movement optimisation

To determine whether males progressively optimised their movement patterns, we analysed return cycles—defined as the number of distinct items visited before returning to a previously visited one— during patrolling and scent-marking over time (Figure 4). If males were optimising their routes, we would expect a gradual increase in return cycles, reflecting more systematic coverage of all available sites and fewer premature revisits. An upper limit would be a return cycle of 12 (i.e., the total number of items), if all sites were visited in perfect sequence without repetition. However, observed return cycles remained consistently below this threshold (Figure 4), indicating that males continued to revisit some locations before fully exploring all available sites.

**Figure 4:**
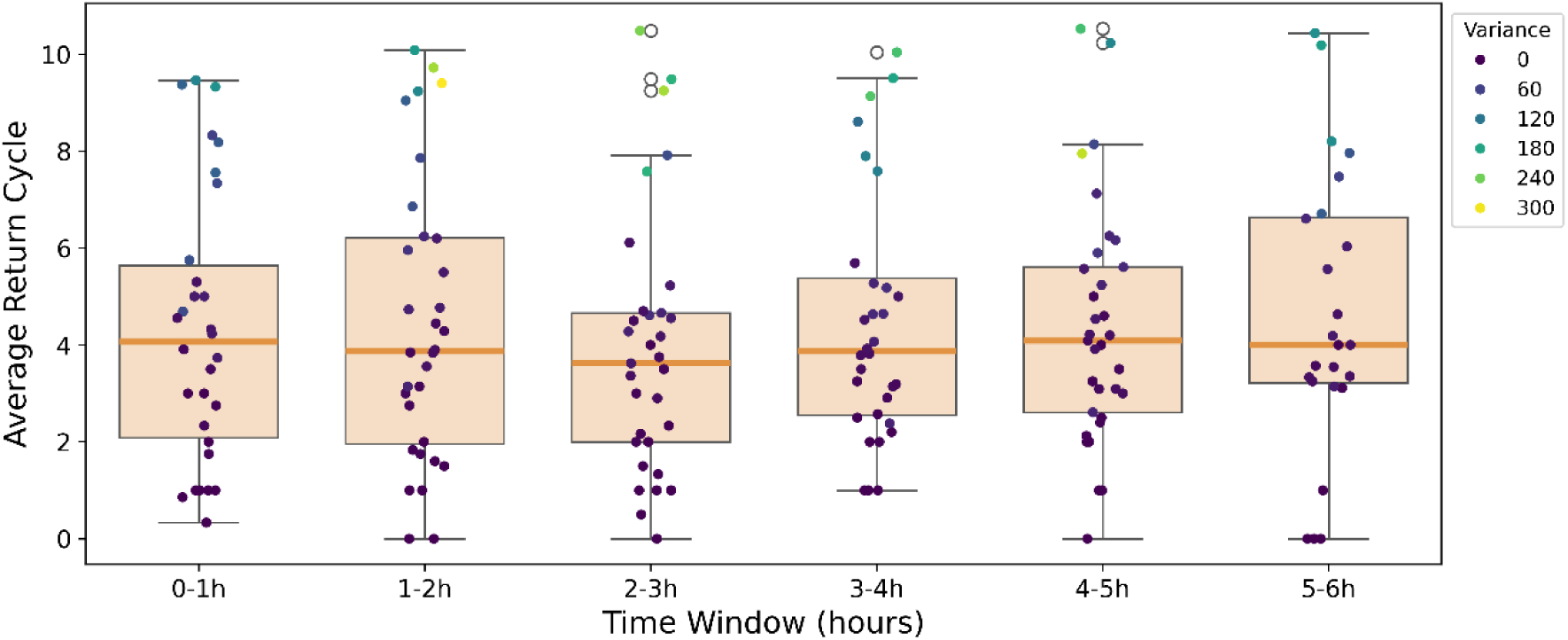
Revisitation patterns in patrolling and scent-marking over time. Boxplots show the average return cycle (number of distinct items visited before returning to a previously visited one) per hour with medians, quartiles, and 10th to 90th percentiles (whiskers). Each dot represents an individual data point, colour-coded by variance in return cycles.

Return cycles remained stable over time (β = 1.44 × 10⁻⁵, SE = 3.39 × 10⁻⁵, t = 0.425, *p* > 0.05), providing no evidence of progressive route optimisation. Variance in return cycles also did not decrease over time, suggesting that movement patterns did not converge toward a more structured or repeatable strategy. While nectar availability had a minor but significant effect (β = 0.1109, SE = 0.0528, z = 2.101, *p* < 0.05), the rate of change was extremely slow (+0.1 per hour), indicating that resource availability had only a minimal influence on revisitation behaviour.

To further examine how experience influences immediate revisitation patterns, we analysed the number of immediate scent-marking, feeding, and patrolling revisits across blocks of visits over time (Figure 5). Here, an immediate revisit was defined as a return to the same location without visiting any other item in between. Males performed fewer immediate scent-marking revisits as they gained experience with the array, suggesting they were less likely to return to the same marking locations in quick succession (β = −0.21, SE = 0.06, z = −3.27, *p* < 0.01). Similarly, immediate feeding revisits declined significantly over time, likely as a response to nectar depletion, prompting males to seek alternative locations (β = −0.12, SE = 0.05, z = −2.42, *p* < 0.05). In contrast, patrolling revisits remained stable, indicating that males did not alter their patrolling revisitation patterns with experience (β = 0.01, SE = 0.02, z = 0.487, *p* > 0.05).

**Figure 5:**
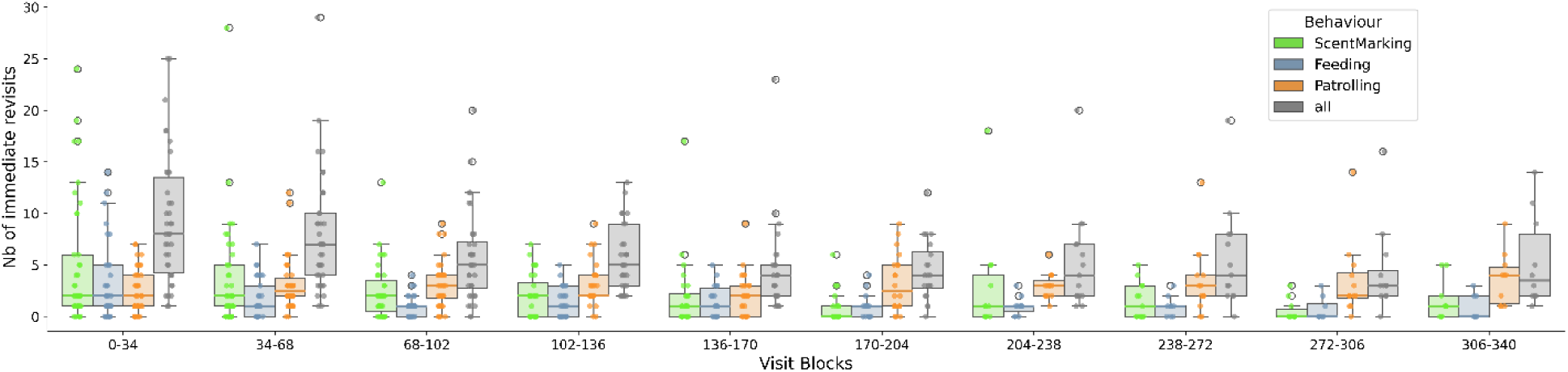
Effect of experience on immediate revisits during scent-marking, feeding, and patrolling. Boxplots show the number of immediate revisits (visits to the same location before all other locations were explored) per block of 34 visits, allowing assessment of changes in revisitation patterns over time. Scent-marking (green), feeding (blue), and patrolling (orange) are represented separately. Boxplots display medians, quartiles, and 10th to 90th percentiles (whiskers), while dots indicate individual data points.

These findings show that while scent-marking and feeding revisits decreased over time—likely due to pheromone accumulation and nectar depletion—patrolling revisits remained stable, reflecting the importance of patrolling in mate-seeking strategies. Overall, males did not systematically optimise their routes, but instead adjusted revisitation patterns based on environmental cues rather than learned, repeatable strategies.

### Route repeatability

To assess structured revisitation patterns, we calculated values of Determinism Index (DET) derived from bee movements and compared them to simulated movement models implementing rigid rules of thumb (Figure 6).

**Figure 6:**
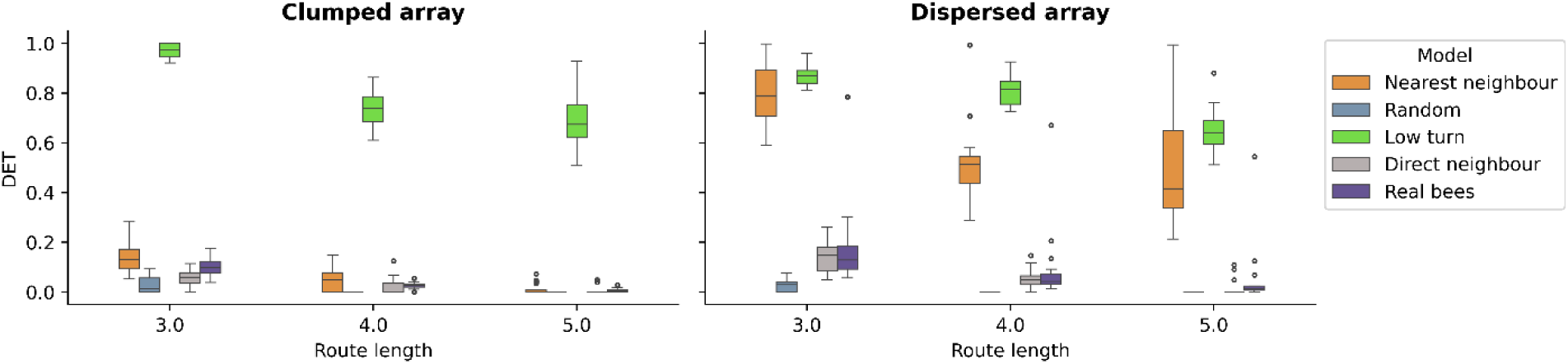
Determinism Index of male bumblebee item visitation sequences in clumped and dispersed arrays, compared to simulated movement models. Boxplots show DET values for real bees and simulated models (nearest neighbour, random, low turn, and direct neighbour) across 3-, 4-, and 5-visit sequences.

In the clumped array, males followed a nearest-neighbour movement pattern, with DET values not significantly different from the nearest-neighbour model (Dunn test after Kruskal–Wallis, *p* > 0.05). However, bumblebee movements significantly differed from random and low-turn models (*p* < 0.001).

In the dispersed array, males’ DET values did not match a single movement rule but were higher than expected for purely random movements. The closest fit was the direct-neighbour model, though DET values were significantly lower than simulated expectations (*p* < 0.05).

Therefore, males in the structured homogenous environment (clumped array) used a predictable nearest-neighbour strategy, while those in the dispersed heterogenous environment display more variable movement patterns, without a clear, consistent strategy.

## DISCUSSION

This study provides new insights into the movement strategies of bumblebee males, showing how spatial resource configuration and nectar availability jointly shape foraging and reproductive behaviours. Unlike bumblebee workers, which tend to optimise their foraging routes for colony provisioning (Lihoreau et al., 2012; Ohashi et al., 2007; Woodgate et al., 2017), males must independently balance energy intake with mate-seeking, leading to movement patterns associated to distinct behavioural trade-offs. Resource distribution had a strong influence on feeding behaviour, while scent-marking and patrolling remained consistent, suggesting a prioritisation of reproductive signalling over energetic optimisation. Nectar availability, somewhat unexpectedly, had minimal effects on these behavioural trade-offs, indicating that mate-seeking investment is maintained even when nectar is scarce. While prior studies have described repeatable individual patrol circuits in males (Frank, 1941; Harano et al., 2018), we found no strong evidence of systematic route formation, possibly reflecting the limited spatial scale of the experimental environment.

### Males maintained reproductive signalling by adjusting foraging behaviour

A key question in male foraging ecology is whether individuals reduce feeding effort to sustain investment in reproductive signalling under varying resource conditions. Our results suggest that males do not reduce signalling behaviours but instead adjust their foraging effort to accommodate energetic constraints. Specifically, scent-marking and patrolling rates remained stable across all nectar availability and spatial configuration treatments, indicating that males maintained mate-seeking behaviour even when the energetic cost of foraging increased.

The dispersed–low nectar condition imposed the highest foraging cost, as reflected by the lowest feeding frequency and total feeding duration. Under these constrained conditions, males did not reduce reproductive investment but instead fed less frequently and for shorter total durations, while extending the duration of individual feeding events when nectar availability allowed. This pattern indicates a context-dependent shift in foraging behaviour that enables males to conserve energy while maintaining consistent signalling effort. Notably, flowers were only refilled once per hour, making it unlikely that longer visits were attempts to wait for replenishment. Instead, longer feeding durations likely reflect more thorough exploitation of the nectar available at the time of arrival. Supporting this, nectar was often observed remaining in the artificial flowers at the end of the day, suggesting that males did not always fully deplete nectar during shorter visits. Rather than a symmetrical trade-off between foraging and reproduction, the data point to a unidirectional adjustment, in which foraging is modulated to preserve reproductive goals.

Revisitation behaviour also varied by behavioural context. Scent-marking and feeding revisits declined over time, likely due to pheromone accumulation and nectar depletion, while patrolling revisits remained stable, underscoring the persistent nature of reproductive effort.

It is important to note that this behavioural stability was observed within the range of nectar conditions tested here. While our nectar delivery rates exceeded average secretion rates for individual wildflowers, they fall within the upper range observed in highly productive plant species and dense floral patches (e.g., *Lamium maculatum*) (Marina et al., 2004). As such, they represent ecologically relevant conditions that bees may encounter in rich environments, rather than extremes. However, under more severe nectar limitation than tested here, males may eventually prioritise feeding over reproductive investment. Future work should explore these lower bounds to determine the robustness and limits of reproductive signalling in *B. terrestris*.

### Spatial heterogeneity and nectar scarcity triggered conservative movement strategies

Male bumblebees adapted their foraging and movement strategies to environmental constraints shaped by both spatial configuration and nectar availability. The clumped array, which was spatially homogeneous with evenly spaced and closely situated resources, supported higher feeding frequency and longer total feeding durations. This likely reflects lower energetic costs, as males predominantly visited items spaced at ∼52 cm—the nearest-neighbour distance—enabling efficient, localised movement.

In contrast, the dispersed array was spatially heterogeneous, with greater variability in inter-item distances and irregular layout. These conditions posed greater energetic challenges. Males in this array fed less frequently and for shorter total durations, especially when nectar was scarce. Movement data showed that under low nectar availability, males exhibited shorter, more consistent transitions and were more likely to revisit previously visited locations, including zero-distance transitions where the same item was visited consecutively. This pattern indicates an energy-conserving foraging strategy, in which males constrained their activity and relied on known sites rather than initiating new exploratory searches.

When nectar availability was high, males in both arrays broadened their foraging range, showing longer and more variable movement transitions—suggesting a shift toward exploratory behaviour aimed at maximising intake from replenished sources. However, this behavioural flexibility does not align with classical optimal foraging models (Pyke, 1984), which predict increased movement when local resource intake is low. Instead, males appeared to prioritise movement efficiency under poor conditions, minimising travel distances and revisiting known sites. This strategy may limit overall nectar intake but likely serves to conserve energy when foraging is most costly.

These findings build on earlier work by Cartar and Real (1997), who showed that worker bumblebees forage more efficiently in spatially structured environments. Our study extends this pattern to male bumblebees, demonstrating that even under different behavioural goals, males benefit from spatial homogeneity through more economical movement and access to resources.

### No evidence of traplining or long-term patrol routes in males

One of the most striking results of this study is the absence of progressive route optimisation by male bumblebees. Unlike workers, males did not refine their foraging paths or reduce variance in their movement patterns over time. Instead, they adopted a flexible, opportunistic approach, frequently revisiting some sites before fully exploring all available ones. This contrasts with trapline foraging observed in workers (Lihoreau et al., 2012; Lihoreau et al., 2013; Ohashi and Thomson, 2009; Reynolds et al., 2013; Saleh and Chittka, 2007), and more broadly with predictions from movement ecology that animals should optimise their routes to improve efficiency (Berger-Tal and Bar-David, 2015).

This difference may reflect both motivational context and spatial scale. Unlike workers, males do not forage to provision the colony, and thus face less pressure to develop efficient, repeatable routes. Their primary selective pressure is reproductive success, which favours sustained signalling over energetic optimisation. While spatial scale may also play a role, our experiment was conducted in a relatively small 3 × 3 m flight cage, where movement costs were low and all sites were accessible with minimal effort. Although traplining has been documented in indoor studies, these typically involved larger arenas (e.g. 5–7 m; Buatois and Lihoreau, 2016; Buatois et al., 2024; Lihoreau et al., 2010; Ohashi et al., 2007), suggesting that both spatial limitations and behavioural priorities contributed to the absence of route learning in our study. Studies comparing bee behaviour in similar arrays of flowers at multiple spatial scales show that bees develop more stable and optimal traplines as distance between flowers increases (Buatois and Lihoreau, 2016).

Early observations described species-typical patrol circuits in male bumblebees (Frank, 1941; Haas, 1946; Haas, 1949). While we observed structured movement patterns, particularly in clumped arrays, we found no evidence of long-term route fidelity or decreasing revisit variance. The discrepancy with earlier observations likely reflects differences in spatial resolution, scale, and ecological context. Future work using harmonic radar (Riley et al., 1996) or other fine-scale tracking in larger scale field conditions could help determine whether male patrol routes become more structured at larger spatial scales.

### Conclusion and future directions

This study offers novel insights into the movement ecology of male bumblebees, showing that they maintain consistent reproductive signalling while flexibly adjusting their foraging strategies to environmental structure. Rather than refining movement paths over time, males appear to adopt a responsive, opportunistic strategy—reducing search effort when necessary, revisiting familiar locations under low resource conditions, and extending feeding durations to conserve energy.

Unlike many other species where males scale reproductive investment to energy reserves (Billings et al., 2019; Dougherty, 2021; Fernlund Isaksson et al., 2022; Meuche and Grafe, 2009), male *B. terrestris* maintain signalling effort even under energetically demanding conditions. This likely reflects strong evolutionary pressure to sustain mate-seeking behaviours despite environmental variability. Instead of cutting back on signalling, males appear to adjust the flexible component of their behavioural budget: foraging. They do so through compensatory strategies, such as increasing feeding duration when nectar is found. In this way, males conserve energy without compromising access to mating opportunities, suggesting that reproductive signalling in this species is behaviourally robust, while foraging effort is the more adaptable component.

Future research should examine whether this behavioural flexibility affects pollination success or mating efficiency, particularly in fragmented or resource-poor landscapes. As landscape structure alters floral distributions, understanding how males navigate complex environments is critical to predicting their ecological roles and reproductive outcomes. Large-scale field studies using tracking systems to record individual flight paths would be especially valuable to assess whether route optimisation emerges at broader spatial scales—or whether flexibility remains the dominant strategy in male movement ecology.

